# *In vivo* MR spectroscopy reflects synapse density in a Huntington’s disease mouse model

**DOI:** 10.1101/2021.10.26.465951

**Authors:** Nicole Zarate, Katherine Gundry, Dahyun Yu, Jordan Casby, Lynn E Eberly, Gülin Öz, Rocio Gomez-Pastor

**Author notes:** Correspondence should be addressed to Rocio Gomez-Pastor, University of Minnesota, 321 Church St. SE, Jackson Hall Room 6-145, Minneapolis, MN 55455. These authors contributed equally to the manuscript (Nicole Zarate, Katherine Gundry).

## Abstract

Striatal medium spiny neurons are highly susceptible in Huntington’s disease (HD), resulting in progressive synaptic perturbations that lead to neuronal dysfunction and death. Non-invasive imaging techniques, such as proton magnetic resonance spectroscopy (^1^H-MRS), are used in HD mouse models and patients with HD to monitor neurochemical changes associated with neuronal health. However, the association between brain neurochemical alterations and synaptic dysregulation is unknown, limiting our ability to monitor potential treatments that may affect synapse function. We conducted *in vivo* longitudinal ^1^H-MRS in the striatum followed by *ex-vivo* analyses of excitatory synapse density of two synaptic circuits disrupted in HD, thalamo-striatal (T-S) and cortico-striatal (C-S) pathways, to assess the relationship between neurochemical alterations and changes in synapse density. We used the zQ175^(Tg/0)^ HD mouse model as well as zQ175 mice lacking one allele of CK2α’(zQ175^(Tg/0)^:CK2α’^(+/−)^), a kinase previously shown to regulate synapse function in HD. Longitudinal analyses of excitatory synapse density showed early and sustained reduction in T-S synapses in zQ175 mice, preceding C-S synapse depletion, which was rescued in zQ175:CK2α’^(+/−)^. Changes in T-S and C-S synapses were accompanied by progressive alterations in numerous neurochemicals between WT and HD mice. Linear regression analyses showed C-S synapse number positively correlated with ^1^H-MRS-measured levels of GABA while T-S synapse number positively correlated with levels of alanine, phosphoethanolamine and lactate, and negatively correlated with total creatine levels.

These associations suggest that these neurochemical concentrations measured by ^1^H-MRS may facilitate monitoring circuit-specific synaptic dysfunction in the zQ175 mouse model and in other HD pre-clinical studies.

**Significance Statement:** The pathogenic events of many neurodegenerative diseases including HD are triggered by reductions in number of synapses. Therefore, *in vivo* measures that reflect synapse number represent a powerful tool to monitor synaptic changes in numerous brain disorders. In this study, we showed that non-invasive *in vivo* ^1^H-MRS reflects excitatory synapse number in the striatum of the zQ175 mouse model of HD. The combination of longitudinal ^1^H-MRS and immunofluorescence synapse detection revealed that distinct neurochemical levels significantly correlated with different striatal glutamatergic synaptic input pathways, suggesting that ^1^H-MRS could distinguish circuit-dependent synapse changes in HD. These results provide potential neurochemical biomarkers to monitor synaptic changes in future pre-clinical trials with HD models.

## INTRODUCTION

Huntington’s disease (HD) is an autosomal dominant, neurodegenerative disorder caused by expansion of a trinucleotide repeat (CAG) in exon 1 of the Huntingtin gene (*HTT)* (MacDonald et al., 1993). The HTT protein is prone to protein misfolding, and its aggregation preferentially affects medium spiny neurons (MSN) of the striatum, causing synaptic perturbations and neuronal death. HD is characterized by progressive motor, cognitive, and psychiatric deficits for which there is no effective therapies. To design therapies that effectively modify disease symptoms it is necessary to identify robust HD biomarkers able to objectively monitor cerebral pathology.

Mouse models of HD and patients with prodromal HD show perturbations in striatal synaptic stability and functional connectivity, respectively, before motor symptom onset or overt neuronal cell death (Raymond et al., 2011; Unschuld et al., 2012). Therefore, having the ability to monitor synaptic stability in the living brain could be a powerful tool to predict disease progression as well as determine the impact of potential therapeutic strategies on synaptic function. However, *in vivo* monitoring of synapses in the human brain is technically challenging. Recently developed Positron Emission Tomography (PET) radioligands targeting various synaptic vesicle proteins can be used to monitor synaptic density in both animals and humans (Finnema et al., 2016; Thomsen et al., 2021). However, the limited availability of these PET tracers and concerns about repeated radiation exposure provide motivation to assess alternative neuroimaging technologies to evaluate synapse density. Magnetic resonance (MR) methods like functional MR imaging (fMRI) and MR spectroscopy (MRS) overcome these concerns but have yet to be evaluated as markers of synaptic density.

MRS has emerged as a useful tool to evaluate neurochemical alterations in HD and other polyQ diseases (Öz et al., 2010; Sturrock et al., 2015; Öz, 2016; Joers et al., 2018). Studies in various mouse models of HD and in patients with HD have highlighted consistent alterations in key neurochemicals that can be associated with neurodegeneration, such as depletion of the neuronal integrity marker *N*-acetylaspartate (NAA) or increased levels of the putative gliosis marker *myo*-inositol (Ins) (Tkác et al., 2007; Heikkinen et al., 2012; Sturrock et al., 2015; Peng et al., 2016). However, the underpinnings of the neurochemical changes that connect them with functional decline is still unknown, limiting our understanding of how potential treatments that may affect synapse function translate into alterations of brain metabolites. Previous studies in patients with different spinocerebellar ataxias (SCAs) presenting different degrees of synapse loss proposed that neurochemical alterations measured by MRS may reflect changes in synaptic function or density (Öz et al., 2011a; Joers et al., 2018). Consistently, a recent MRS study in aged zQ175 mice showed that key striatal neurochemical alterations paralleled changes in T-S synapses (Yu et al., 2020), highlighting the potential use of MRS as a tool to monitor synapse density. However, the relationship between neurochemical alterations and synaptic density has not been directly assessed thus far.

In the present study we conducted *in vivo* longitudinal high field proton (^1^H) MRS in the striatum of the heterozygous zQ175 HD mouse model followed by *ex-vivo* analyses of excitatory synapse densities of two major, differentially altered striatal synaptic circuits in HD; cortico-striatal (C-S) and thalamo-striatal (T-S). We also studied zQ175 mice lacking one allele of CK2α’, a kinase involved in synapse stability in HD, the haploinsufficiency of which differentially altered T-S and C-S circuitry and ameliorated long-term HD-like symptoms (Gomez-Pastor et al., 2017; Yu et al., 2020). We found that T-S synaptic density was significantly correlated with alterations in alanine (Ala), lactate (Lac), phosphoethanolamine (PE), and total creatine (tCr) while C-S synapse number significantly correlated with gamma-aminobutyric acid (GABA). We propose that GABA, Ala, Lac, PE, and tCr alterations could be used as surrogate biomarkers to monitor circuit-dependent synapse dysfunction during HD in zQ175 mice. Furthermore, these results support using MRS as a technique in future HD pre-clinical studies to monitor treatment efficacy in altering excitatory synaptic function in the striatum.

## METHODS

### Experimental Design

A cohort of mice (WT n=13, 8 females, 5 males; zQ175 n=16, 7 females, 9 males; and zQ175:CK2α’^(+/−)^ n=16, 11 females, 5 males) were scanned at 3 months of age using a 9.4T scanner, after which a subset of the animals (n=4 per genotype) were sacrificed for synapse density analyses (n=3 per genotype were used). One additional zQ175:CK2α’^(+/−)^ mouse was added to the cohort after the 3 months scan. Remaining animals were aged to 6 months at which time they were again scanned (WT n=9, 7 females, 2 males; zQ175 n=12, 5 females, 7 males; and zQ175:CK2α’^(+/−)^ n=13, 9 females, 4 males), and another subset (n=4 per genotype) sacrificed for synapse density analyses. Two zQ175:CK2α’^(+/−)^ mice were harvested prior to being scanned at 12 months due to health issues. The remaining animals (WT n=5 (3 females, 2 males), zQ175 n=8 (4 females, 4 males), and zQ175:CK2α’^(+/−)^ n=7 (4 females, 3 males) were scanned at 12 months followed by striatal synapse density analyses (only n=3 (2 females, 1 male) were used for zQ175:CK2α’^(+/−)^ synapse analysis). CK2α’^(+/−)^ control mice were not included in the experimental design since they did not show significant changes in either T-S or C-S circuits compared to WT mice (Gomez-Pastor et al., 2017). We selected these three time points based on progression of motor symptoms (Heikkinen et al., 2012; Menalled et al., 2012): 3 months (pre-symptomatic), 6 months (early symptomatic) and 12 months (symptomatic). Longitudinal neuroimaging provided information for how neurochemical levels changed within individual mice over time. A diagram illustrating the experimental design is shown in **Fig. 1.**

**Figure 1.**
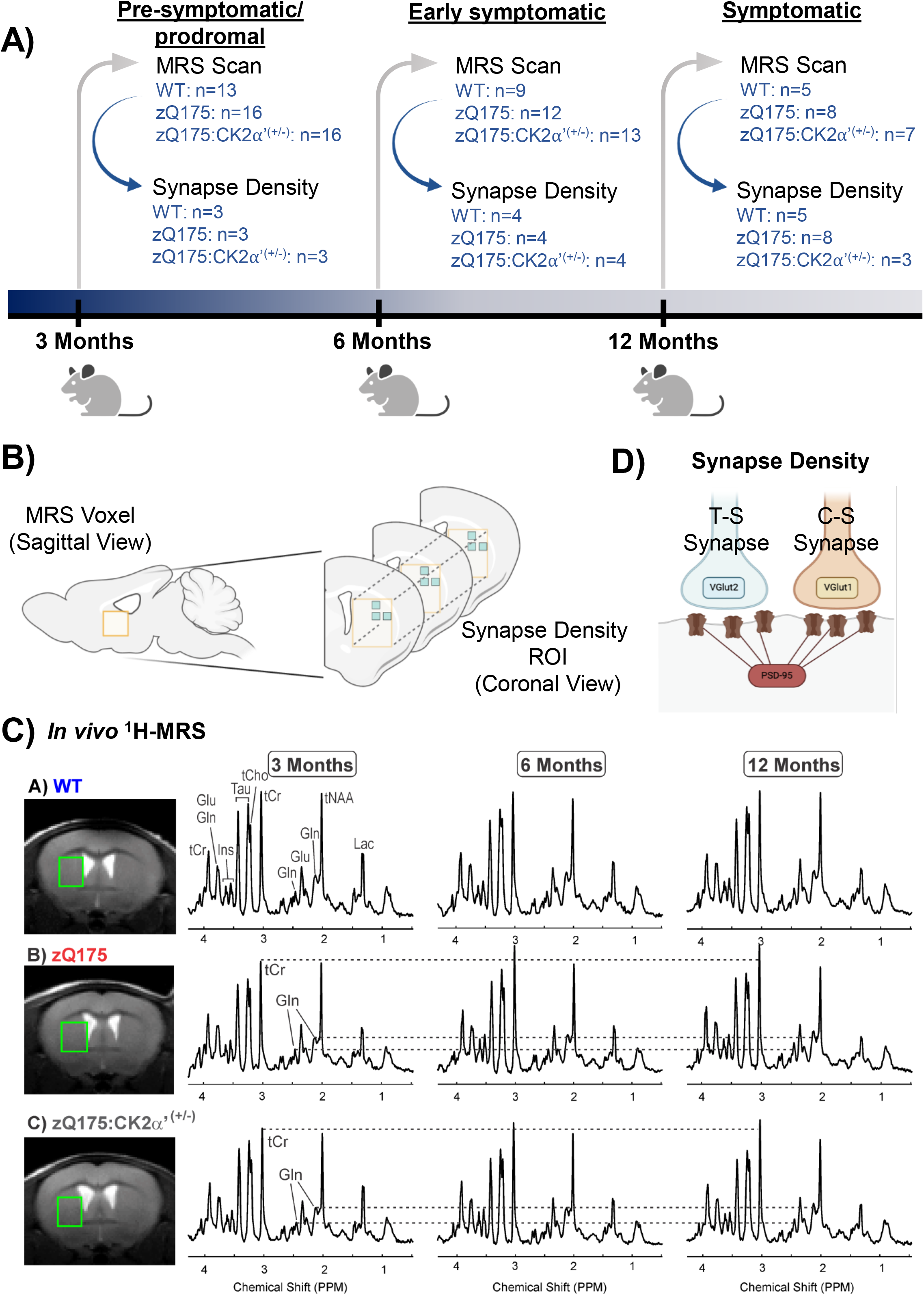
Diagram of experimental design and timeline. **A**) Genotypes and number of animals used for each analysis are annotated. A cohort of WT, zQ175 and zQ175:CK2α’^(+/−)^ mice were scanned longitudinally on a 9.4T magnet (MRS) and a subset of mice were sacrificed for synapse density analyses by immunohistochemistry at each time point. **B**) Diagrams indicate the brain region covered by MRS and synapse analysis. Yellow box indicates voxel range (Volume = 8.2 μl), blue box indicates synapse density ROI within the voxel range. Synapse density was analyzed as described in the Methods and Materials section by conducting PSD-95 and VGlut1 or VGlut2 co-localization analyses from at least n=6-9 images corresponding to n=3 coronal sections. Graphics were created with Biorender.com **C**) Localized proton MR spectra measured from the mouse dorsolateral striatum in WT, zQ175^(Tg/0)^ and zQ175^(Tg/0)^:CK2α’^(+/−)^ mice at 3, 6, and 12 months old. The volume of interest is shown on T2-weighted images and alterations in neurochemicals visible in the spectra are shown.

### Animal Preparation for and Monitoring during MR Scanning

All experiments were performed according to procedures approved by the University of Minnesota Institutional Animal Care and Use Committee. Animals were induced with 3% isoflurane in a 1:1 mixture of O_2_:N_2_O. Mice were secured in a custom-built mouse holder and physiological status was monitored (SA Instruments) and recorded. Anesthesia was maintained with 1.5-2% isoflurane to achieve a respiration rate of 70-100 breaths per minute. Body temperature was maintained at 36-37°C with a circulating warm water system and a heating fan controlled by feedback received from a fiber-optic rectal thermometer. The scan session was approximately 50 minutes for each animal.

### MR Protocol

All experiments were performed on a 9.4T/31 cm scanner (Agilent), as described previously (Öz et al., 2015; Friedrich et al., 2018). A quadrature surface radio frequency (RF) coil with two geometrically decoupled single turn coils (14 mm diameter) was used as the MR transceiver. Following positioning of the mouse in the magnet, coronal and sagittal multislice images were obtained using a rapid acquisition with relaxation enhancement (RARE) sequence (Hennig et al., 1986) [repetition time (TR)= 4 s, echo train length= 8, echo time (TE)= 60 ms, slice thickness= 1 mm, 7 slices]. The volume of interest (VOI) was centered on the striatum (8.2 μl, 1.7 × 2.0 × 2.4 mm^3^). All first- and second-order shims were adjusted using FASTMAP with echo-planar readout (Gruetter and Tkác, 2000). Localized ^1^H MR spectra were acquired with a short-echo localization by adiabatic selective refocusing (LASER) sequence [TE= 15 ms, TR= 5 s, 256 transients] (Garwood and DelaBarre, 2001) combined with VAPOR (variable power RF pulses with optimized relaxation delays) water suppression (Tkác et al., 1999). Spectra were saved as single scans. Unsuppressed water spectra were acquired from the same VOI for metabolite quantification.

### Metabolite quantification

Single shots were eddy current, frequency, and phase corrected using MRspa software (http://www.cmrr.umn.edu/downloads/mrspa/) before averaging. The contributions of individual metabolites to the averaged spectra were quantified using LCModel (Provencher, 1993) as described previously (Friedrich et al., 2018). The following metabolites were included in the basis set: alanine (Ala), ascorbate/vitamin C (Asc), aspartate, glycerophosphocholine (GPC), phosphocholine (PCho), creatine (Cr), phosphocreatine (PCr), gamma-aminobutyric acid (GABA), glucose (Glc), glutamine (Gln), glutamate (Glu), glutathione, glycine, *myo*-inositol (Ins), lactate (Lac), *N*-acetylaspartate (NAA), *N*-acetylaspartylglutamate (NAAG), phosphoethanolamine (PE), taurine (Tau), and macromolecules (MM). The MM spectra were experimentally obtained from a VOI that covered the striatum using an inversion recovery technique [VOI = 4.7 × 2.1 × 2.7 mm^3^, TE = 15 ms, TR = 2.0 s, inversion time (TIR) = 675 ms, 400 transients, N = 2]. The model metabolite spectra were generated using density matrix simulations (Govindaraju et al., 2000) with the MATLAB software (MathWorks) based on previously reported chemical shifts and coupling constants (Govindaraju et al., 2000; Tkác, 2008). Concentrations with mean Cramér-Rao lower bounds (CRLB) ≤20% in any of the 3 groups were reported (Friedrich et al., 2018). If the correlation between two metabolites was consistently high (correlation coefficient r <−0.7), their sum was reported rather than the individual values (Friedrich et al., 2018), this was the case for total NAA (tNAA= NAA+NAAG), total Cr (tCr= Cr+PCr) and total Cho (tCho= GPC+PCho). In addition, Glc + Tau was reported in addition to the separate Glc and Tau concentrations because the individual concentrations did not always meet the mean CRLB ≤20% criterion and Glc and Tau have similar spectral patterns.

### Synapse density quantification

Two to three independent coronal brain sections were used for each mouse, containing the dorsal striatum (bregma 0.5–1.1 mm) and were stained with presynaptic VGlut1 (Millipore AB5905, 1:500) or VGlut2 (Millipore AB2251-I1:1000) and postsynaptic PSD95 (Thermofisher 51-6900, 1:500) markers as described previously (Ippolito and Eroglu, 2010; Mckinstry et al., 2014; Gomez-Pastor et al., 2017). Secondary antibodies used were goat anti-guinea pig Alexa 488 (VGlut1/2, Invitrogen A-11073, 1:200) and goat anti-rabbit Alex 594 (PSD-95, Invitrogen A-11012, 1:200). At least three mice for each genotype; WT, zQ175 and zQ175:CK2α’^(+/−)^ were evaluated. The confocal scans (optical section depth 0.34 mm, 15 sections per scan) of the synaptic zone in the dorsal striatum were performed at 60X magnification on an Olympus FV1000 confocal laser-scanning microscope. Maximum projections of three consecutive optical sections were generated. The Puncta Analyzer Plugin for ImageJ was used to count co-localized synaptic puncta in a blinded fashion. This assay takes the advantage of the fact that presynaptic and postsynaptic proteins reside in separate cell compartments (axons and dendrites, respectively), and they would appear to co-localize at synapses because of their close proximity. The number of animals used in this analysis was 3 per genotype (3 month timepoint), 4 per genotype (6 month timepoint), and 5-6 WT, 7-8 zQ175, and 3 zQ175:CK2α’^(+/−)^ (12 month timepoint).

### Data Analysis

All statistics were conducted using GraphPad Prism 9 software. Group averages between genotypes were compared at each time point using one-way ANOVA for MRS and synapse densities with Tukey’s post-hoc test to adjust for multiple comparisons. Significant group differences for synapse density were calculated using n=6-9 images per animal, 3 animals minimum per genotype, resulting in a minimum of 18 data points per genotype for statistical analyses. Only averages for each mouse are shown. Pearson correlation analyses were run separately for each neurochemical against synapse number. Holm-Šídák adjustment of correlation p-values was used to correct for the multiple testing of the many neurochemicals together with metabolite sums and metabolite ratio. A full report of results for MRS ANOVAs are in Table 1, with synapse analyses in Table 2, and correlation analyses in Table 3. Fisher’s exact test was conducted to evaluate sex differences for all metabolites (none were found). Error bars always represent mean ± standard deviation (SD).

**Table 1.**
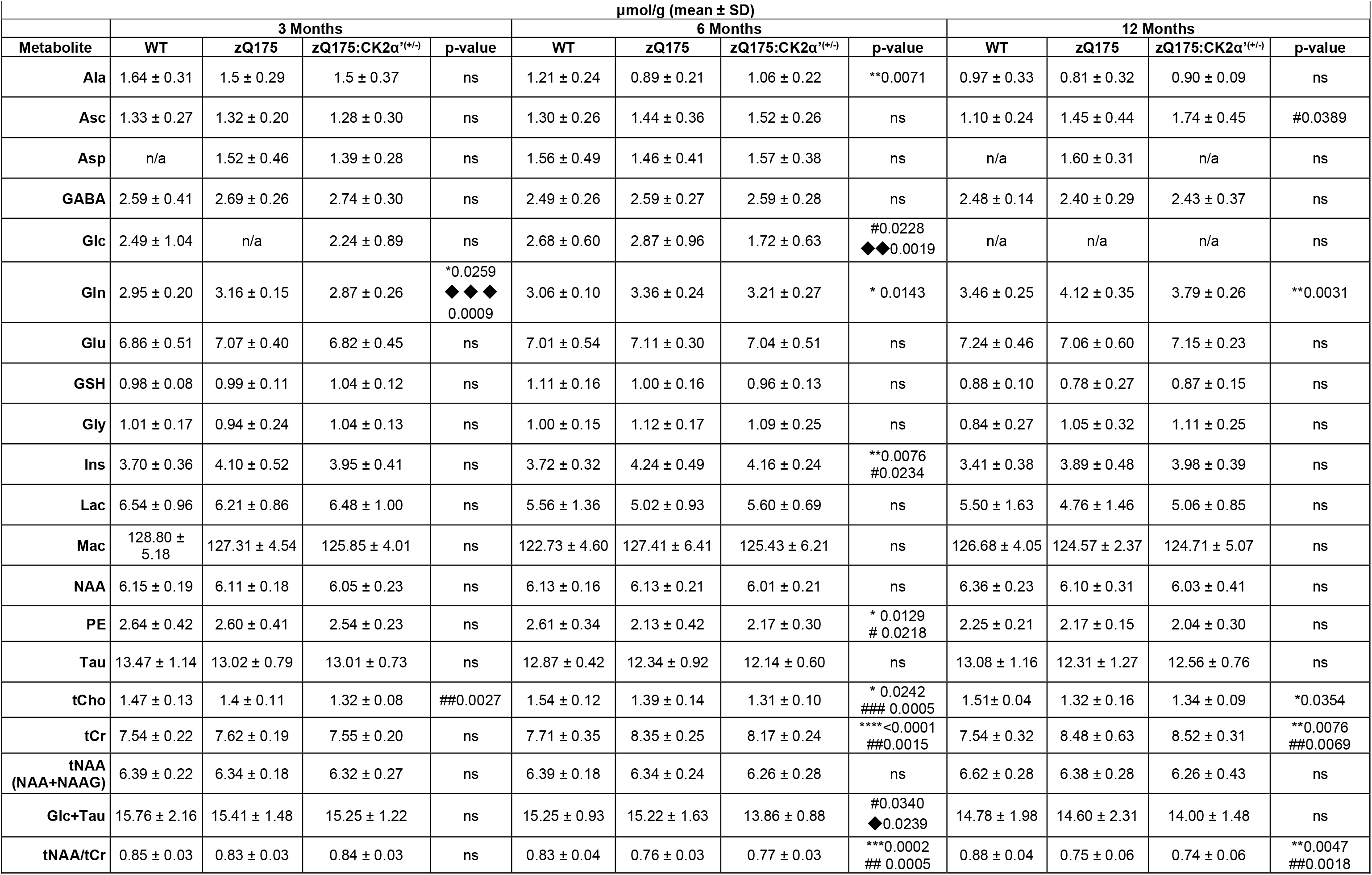
ANOVA results for neurochemical concentrations measured by ^1^H-MRS. Neurochemical concentrations in WT, zQ175, and zQ175:CK2α’^(+/−)^ mice over time. Values represented as mean ± SD. One-way ANOVA with Tukey-adjusted p-values for those <0.05. * indicates WT vs. zQ175, # indicates WT vs. zQ175:CK2α’^(+/−)^, and ⧫ indicates zQ175 vs. zQ175:CK2α’^(+/−)^.

**Table 2.** Excitatory synapse density ANOVA results. Mean synapse density (relative to WT) between genotypes and across time points. Error presented as ± SD. One-way ANOVA with Tukey-adjusted p-values.

| **C-S Synapses** |  |  |  |  |  |  |
| --- | --- | --- | --- | --- | --- | --- |
|  | 3 months |  | 6 Months |  | 12 Months |  |
|  | Mean<br>(% of WT) | Tukey-Adjusted<br>p-value | Mean<br>(% of WT) | Tukey-Adjusted<br>p-value | Mean<br>(% of WT) | Tukey-Adjusted<br>p-value |
| WT vs zQ175 | 100.00 ± 32.83<br>vs<br>114.4 ± 24.68 | 0.5769 | 100.00 ± 22.13<br>vs<br>85.61 ± 21.01 | 0.1976 | 100.00 ± 30.19<br>vs<br>59.35 ± 2.073 | <0.0001 |
| WT vs<br>zQ175:CK2α<sup>(+/-)</sup> | 100.00 ± 32.83<br>vs<br>135.8 ± 36.17 | 0.0031 | 100.00 ± 22.13<br>vs<br>83.14 ± 16.39 | 0.0425 | 100.00 ± 30.19<br>vs<br>69.31 ± 22.19 | <0.0001 |
| zQ175 vs<br>zQ175:CK2α<sup>(+/-)</sup> | 114.4 ± 24.68<br>vs<br>135.8 ± 36.17 | 0.0628 | 85.61 ± 21.01<br>vs<br>83.14 ± 16.39 | 0.7661 | 59.35 ± 2.073<br>vs<br>69.31 ± 22.19 | 0.4883 |
| **T-S Synapses** |  |  |  |  |  |  |
|  | 3 months |  | 6 Months |  | 12 Months |  |
|  | Mean<br>(% of WT) | Tukey-Adjusted<br>p-value | Mean<br>(% of WT) | Tukey-Adjusted<br>p-value | Mean<br>(% of WT) | Tukey-Adjusted<br>p-value |
| WT vs zQ175 | 100.00 ± 16.01<br>vs<br>79.61 ± 16.02 | 0.0134 | 100.00 ± 32.97<br>vs<br>62.17 ± 35.45 | 0.013 | 100.00 ± 30.23<br>vs<br>72.73 ± 22.58 | 0.0062 |
| WT vs<br>zQ175:CK2α<sup>(+/-)</sup> | 100.00 ± 16.01<br>vs<br>101.0 ± 10.54 | 0.9778 | 100.00 ± 32.97<br>vs<br>108.0 ± 58.73 | 0.4771 | 100.00 ± 30.23<br>vs<br>88.32 ± 65.25 | 0.5932 |
| zQ175 vs<br>zQ175:CK2α<sup>(+/-)</sup> | 79.61 ± 16.02<br>vs<br>101.0 ± 10.54 | 0.0083 | 62.17 ± 35.45<br>vs<br>108.0 ± 58.73 | 0.0004 | 72.73 ± 22.58<br>vs<br>88.32 ± 65.25 | 0.2901 |

**Table 3.**
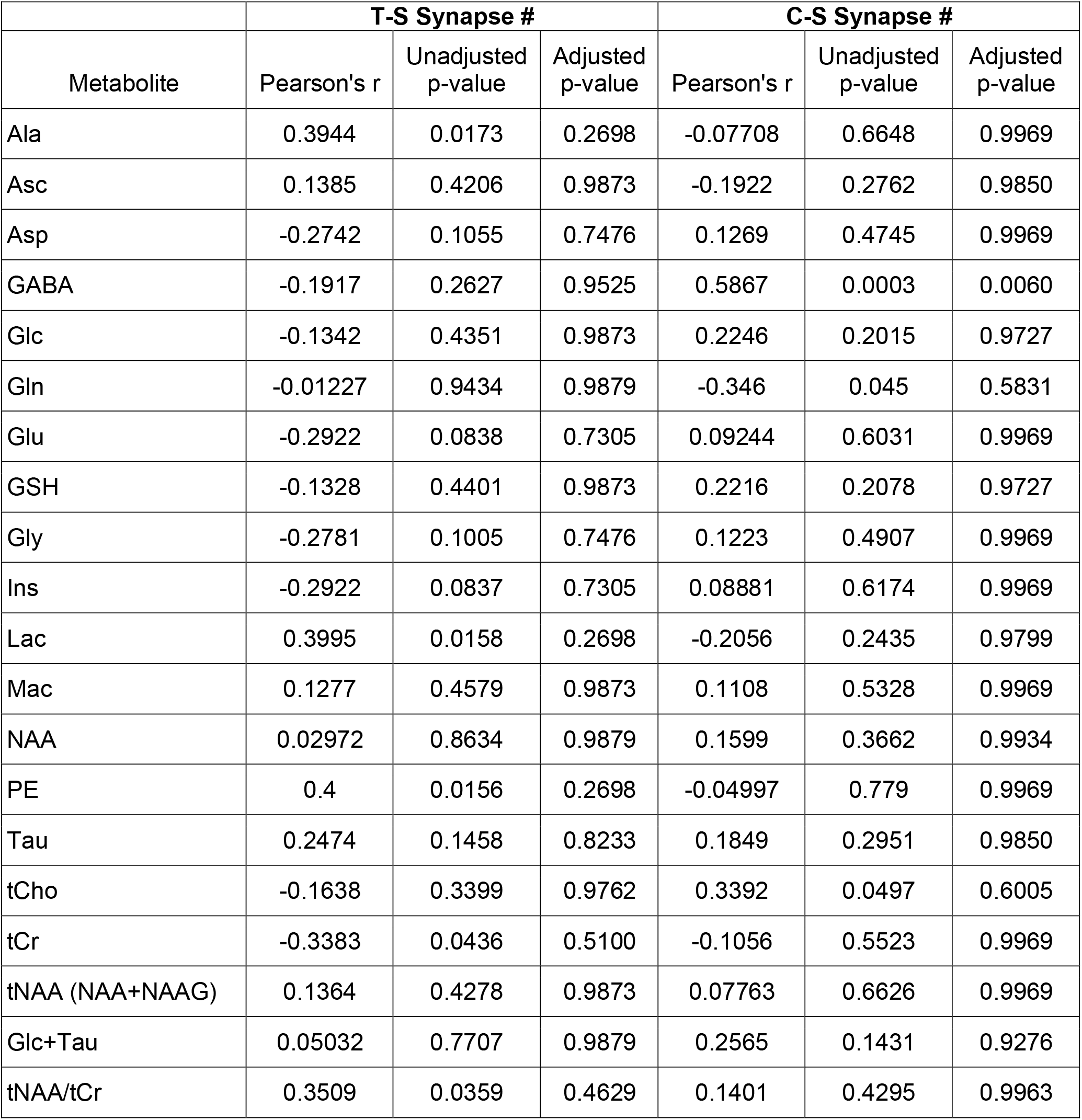
Correlation analyses between synapse number and metabolite concentration. Pearson coefficient and unadjusted p-value calculated from correlation analyses between all neurochemicals and synapse # (C-S and T-S correlations run separately). Holm-Šídák multiple comparison post-hoc test was run for C-S and T-S separately, represented by adjusted p-values. All ages and genotypes are pooled for each analysis corresponding to WT n=12, zQ175 n=15 and zQ175:CK2α’^(+/−)^ n=9 from all three time points (3, 6 and 12 months).

| Metabolite | T-S Synapse # |  |  | C-S Synapse # |  |  |
| --- | --- | --- | --- | --- | --- | --- |
|  | Pearson's r | Unadjusted p-value | Adjusted p-value | Pearson's r | Unadjusted p-value | Adjusted p-value |
| Ala | 0.3944 | 0.0173 | 0.2698 | -0.07708 | 0.6648 | 0.9969 |
| Asc | 0.1385 | 0.4206 | 0.9873 | -0.1922 | 0.2762 | 0.9850 |
| Asp | -0.2742 | 0.1055 | 0.7476 | 0.1269 | 0.4745 | 0.9969 |
| GABA | -0.1917 | 0.2627 | 0.9525 | 0.5867 | 0.0003 | 0.0060 |
| Glc | -0.1342 | 0.4351 | 0.9873 | 0.2246 | 0.2015 | 0.9727 |
| Gln | -0.01227 | 0.9434 | 0.9879 | -0.346 | 0.045 | 0.5831 |
| Glu | -0.2922 | 0.0838 | 0.7305 | 0.09244 | 0.6031 | 0.9969 |
| GSH | -0.1328 | 0.4401 | 0.9873 | 0.2216 | 0.2078 | 0.9727 |
| Gly | -0.2781 | 0.1005 | 0.7476 | 0.1223 | 0.4907 | 0.9969 |
| Ins | -0.2922 | 0.0837 | 0.7305 | 0.08881 | 0.6174 | 0.9969 |
| Lac | 0.3995 | 0.0158 | 0.2698 | -0.2056 | 0.2435 | 0.9799 |
| Mac | 0.1277 | 0.4579 | 0.9873 | 0.1108 | 0.5328 | 0.9969 |
| NAA | 0.02972 | 0.8634 | 0.9879 | 0.1599 | 0.3662 | 0.9934 |
| PE | 0.4 | 0.0156 | 0.2698 | -0.04997 | 0.779 | 0.9969 |
| Tau | 0.2474 | 0.1458 | 0.8233 | 0.1849 | 0.2951 | 0.9850 |
| tCho | -0.1638 | 0.3399 | 0.9762 | 0.3392 | 0.0497 | 0.6005 |
| tCr | -0.3383 | 0.0436 | 0.5100 | -0.1056 | 0.5523 | 0.9969 |
| tNAA (NAA+NAAG) | 0.1364 | 0.4278 | 0.9873 | 0.07763 | 0.6626 | 0.9969 |
| Glc+Tau | 0.05032 | 0.7707 | 0.9879 | 0.2565 | 0.1431 | 0.9276 |
| tNAA/tCr | 0.3509 | 0.0359 | 0.4629 | 0.1401 | 0.4295 | 0.9963 |

## RESULTS

### Longitudinal *in vivo* ^1^H-MRS captures genotype and age-specific striatal neurochemical changes in the zQ175 HD mouse model

MRS studies have been widely applied to evaluate neurodegeneration in several mouse models of HD and patients with HD (Tkác et al., 2007; Sturrock et al., 2010, 2015; Heikkinen et al., 2012; Peng et al., 2016). The use of mouse models allows for parallel characterization of neurochemical concentrations and neuropathological features, an essential aspect of understanding how different neurochemical abnormalities relate to pathology. We chose the heterozygous knock-in zQ175 HD mouse model (zQ175^Tg/0^) for its slow disease progression and its ability to recapitulate multiple HD-like features observed in patients with HD, such as HTT aggregation, progressive synaptic deficits, transcriptional dysregulation, gliosis, weight loss, and motor and cognitive impairment (Heikkinen et al., 2012; Menalled et al., 2012; Gomez-Pastor et al., 2017).

We utilized *in vivo* ^1^H-MRS to measure neurochemicals in the striatum of WT, zQ175, and zQ175:CK2α’^(+/−)^ mice in a longitudinal manner and in parallel with *ex vivo* synapse density analyses to obtain simultaneous information for potential neurochemical and synapse alterations across different genotypes and time points (**Fig. 1**). A total of 19 neurochemicals, including 4 sums, were quantified reliably in the dorsolateral striatum (**Table 1**). In addition, tNAA/tCr ratio was computed as it is frequently utilized as a marker of neuronal viability and to enable comparison of the current results with prior literature. Seven neurochemicals (Ala, Asc, Gln, Ins, PE, tCho, and tCr) and tNAA/tCr showed significant group differences primarily at 6 and 12 months of age (**Fig. 2)**. Ala, Asc, Ins, and PE (**Fig. 2A, B, C, D)** showed a decrease with age in WT mice while Gln (**Fig. 2E**) increased. Similar age-dependent alterations in brain neurochemicals have also been reported in C57BL/6 mice and in human studies comparing young and middle-aged subjects (Grachev and Apkarian, 2001; Duarte et al., 2014), indicating these alterations are common brain modifications during aging.

**Figure 2.**
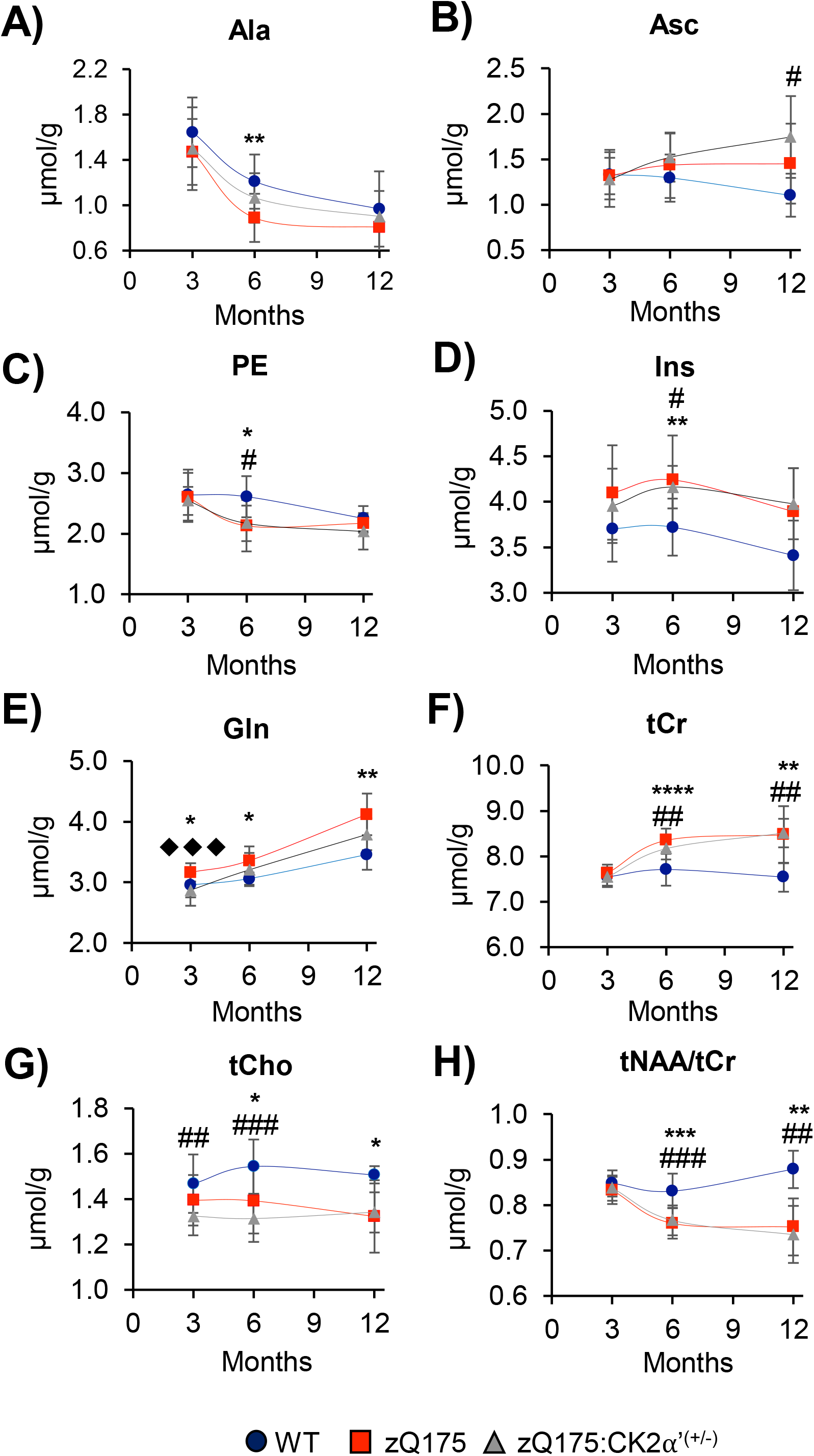
Age and genotype-dependent neurochemical alterations in the mouse dorsolateral striatum. Longitudinal metabolite profile in WT, zQ175, and zQ175:CK2α’^(+/−)^ mice. Only neurochemicals that showed significant differences between genotypes are shown (see Table 1). Error bars represent mean ± SD. One-way ANOVA with Tukey’s post-hoc test. *p<0.05 WT vs. zQ175, # p<0.05 WT vs. zQ175:CK2α’^(+/−)^, ⧫ p<0.05 zQ175 vs. zQ175:CK2α’^(+/−)^. Abbreviations can be found in the methods section. For 3 month WT n=13, zQ175 n= 16, zQ175:CK2α’^(+/−)^ n=16. 6 month WT n=9, zQ175 n= 12, zQ175:CK2α’^(+/−)^ n=13. 12 month WT n=5, zQ175 n= 8, zQ175:CK2α’^(+/−)^ n=7.

The levels of several neurochemicals in zQ175 mice significantly differed compared to WT at multiple time points. Gln was significantly higher in zQ175 vs. WT starting at 3 months and increased even further at 6 and 12 months (**Fig. 2C**). tCr was significantly higher and tCho lower in zQ175 mice vs. WT at 6 and 12 months (**Fig. 2F, G**) coinciding with previously reported worsening motor and cognitive symptoms. On the other hand, no significant differences in tNAA were observed between WT and zQ175 mice while a higher Ins level in zQ175 mice reached statistical significance only at 6 months (**Fig. 2D, Table 1**), recapitulating what was previously reported for this mouse model (Heikkinen et al., 2012). When tNAA was referenced to tCr, which is often used as an internal concentration reference, we found a significantly lower (tNAA/tCr) ratio in zQ175 vs. WT mice at 6 and 12 months (**Fig. 2H**). This ratio was relatively stable in WT mice over time, but it showed an age-dependent decrease in zQ175 mice. However, this alteration most likely reflects changes in tCr. Lower NAA and higher Ins, which are considered markers of neurodegeneration, were previously reported in more severe HD mouse models such as the R6/2 (12 weeks) (Tkác et al., 2007), homozygous zQ175^(Tg/Tg)^ (12 months) (Peng et al., 2016) and in older zQ175 ^Tg/0^ (22 months) (Tkác et al., 2007; Heikkinen et al., 2012; Peng et al., 2016; Yu et al., 2020). Therefore, these data suggests that tNAA and Ins may reflect substantial neurodegeneration in advanced disease. On the other hand, changes in Gln, tCr, tCho and tNAA/tCr in HD mice coincided with symptom onset and progression for this HD mouse model.

Given the previously described positive effects of CK2α’ haploinsufficiency in ameliorating HD-like symptoms (Gomez-Pastor et al., 2017; Yu et al., 2020), we expected to see a number of neurochemicals that were significantly different between zQ175 and zQ175:CK2α’^(+/−)^ mice. Interestingly, we only observed significant differences between these two genotypes at 3 months for Gln (**Fig. 2E**), while no significant differences were observed between WT and zQ175:CK2α’^(+/−)^ mice at the later time points, indicating that the characteristic increase in Gln in zQ175 could be delayed as a consequence of manipulating CK2α’ levels. zQ175:CK2α’^(+/−)^ mice have previously shown decreased HTT aggregation in the striatum, increased excitatory synaptic transmission and improved motor coordination compared to zQ175 (Yu et al., 2020). The early lower Gln level in zQ175:CK2α’^(+/−)^ vs. zQ175 mice may therefore reflect those pathological features.

### Loss of thalamo-striatal excitatory synapses precedes cortico-striatal synapse loss and is sustained over time

MSNs receive excitatory glutamatergic input from the cortex and intralaminar nuclei of the thalamus and both D1 (direct pathway) and D2-type MSNs (indirect pathway) rely equally on these excitatory inputs (Smith et al., 2009; Huerta-Ocampo et al., 2014). Dysfunction in the cortico-striatal (C-S) pathway has been widely reported in HD, showing disruptions in this circuitry occur prior to MSN death thereby suggesting they are the leading cause for neuronal dysfunction (Cepeda et al., 2007; Raymond et al., 2011; Unschuld et al., 2012). However, recent studies in cell and mouse models of HD showed that thalamic input is also disrupted in HD and occurs before C-S circuit pathology or onset of motor symptoms (Cepeda et al., 2007; Mckinstry et al., 2014; Kolodziejczyk and Raymond, 2016; Gomez-Pastor et al., 2017), indicating that the thalamo-striatal (T-S) excitatory circuit may play a more prominent role in early HD pathogenesis than previously considered.

To study whether specific neurochemical alterations relate to changes in synapse density, we first determined changes in excitatory synapse density in WT, zQ175, and zQ175:CK2α’^(+/−)^ mice, looking at both C-S and T-S circuitries during disease progression (**Fig. 3A**). Synapse density was quantified using immunofluorescent colocalization of the presynaptic markers VGlut1 (Vesicular glutamate transporter 1: specific marker for cortical input) and VGlut2 (Vesicular glutamate transporter 2: specific marker for thalamic input) (Fujiyama et al., 2004; Huerta-Ocampo et al., 2014) and the post-synaptic marker PSD-95 (post-synaptic density protein 95) in the dorsal striatum of 3, 6, and 12 month old animals (**Fig. 3B**). C-S synapses were initially higher in zQ175:CK2α’^(+/−)^ mice compared to WT, however by 6 months this trend had reversed and by 12 months had become significantly lower in both zQ175 and zQ175:CK2α’^(+/−)^ mice compared to WT (**Fig. 3C, Table 2**). In contrast, levels of T-S synapses in zQ175 mice were significantly reduced compared to WT at all 3 time points while these decreases were significantly rescued in zQ175:CK2α’^(+/−)^ mice at 3 and 6 months (**Fig. 3D, Table 2**). These results validated previous observations in zQ175:CK2α’^(+/−)^ mice, showing ameliorated T-S synapse loss and long-term HD-like symptoms with limited impact on C-S synapse density (Gomez-Pastor et al., 2017). The lack of statistically significant rescue at 12 months between zQ175 and zQ175:CK2α’^(+/−)^ suggests that CK2α’ haploinsufficiency delays the onset of T-S synapse loss but does not prevent progression. Taken together, these data further demonstrate that there are circuit-specific changes in striatal excitatory synapse dysfunction in HD that, in the T-S pathway, are in part ameliorated by reducing levels of CK2α’.

**Figure 3.**
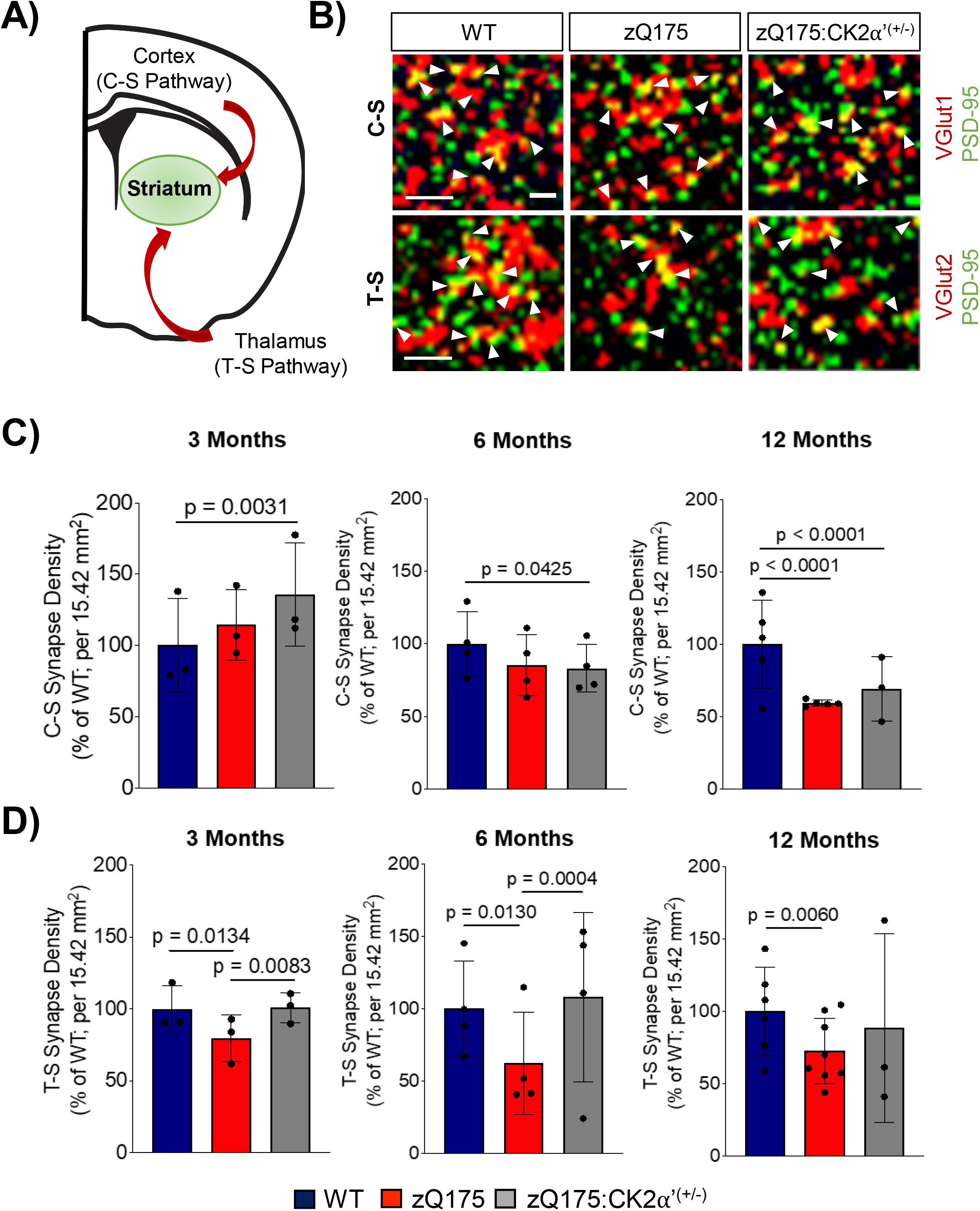
Onset of thalamo-striatal (T-S) synapse loss precedes cortico-striatal synapse deficits in zQ175 mice and it is delayed in zQ175:CK2α’^(+/−)^ mice. **A**) Diagram of the striatal excitatory circuitry. **B**) Colocalization (white arrows) of pre-synaptic (VGlut1/2) and post-synaptic (PSD-95) markers in WT, zQ175, and zQ175:CK2α’^(+/−)^ mice. Representative images from 6 months. Scale bar: 5 μm. **C-D**) Quantification of C-S and T-S synapses respectively at 3, 6, and 12 months old between genotypes. At each time point, WT n= 3-6, zQ175 n= 3-8, zQ175:CK2α’^(+/−)^ n=3-4. Significant group differences were calculated using n=6-9 images per animal resulting in a minimum of 18 data points per genotype for statistical analyses. Only averages for each individual mouse are shown. Error bars represent mean ± SD. *p<0.05, **p<0.01, ***p<0.001, ****p<0.0001, one-way ANOVA with Tukey’s post-hoc test.

### Changes in striatal neurochemical levels correlate with circuit-dependent changes in synapse density

Given the alterations in neurochemical levels and changes in excitatory synapse densities between the different genotypes, we wanted to determine if these two measures were significantly correlated with each other, potentially allowing us to identify surrogate biomarkers for synapse loss in HD. We performed correlation analyses across all mice used in this study (n=36). Synapse number for each mouse analyzed at one of three time points (3, 6 and 12 months) was used as the predictor variable in the analysis and neurochemical levels at the same time points as the response variable (**Fig. 4**, **Table 3**). Regressions were run separately to examine each synaptic circuit, T-S and C-S, and their correlation with each neurochemical level. When analyzed in this manner, we found a significant positive correlation (p <0.05) between C-S synapse number and GABA (**Fig. 4A, B**). Statistical significance of this correlation was maintained even after conservative multiple testing adjustment of p-values across all the metabolites. Gln and tCho had statistically significant un-adjusted p-values but only modestly sized correlations. GABA is known for its role as the primary inhibitory neurotransmitter in the adult brain and dysfunction in GABAergic signaling has been implicated in HD mouse models as well as patients with HD (Hsu et al., 2018). It is important to note that despite the positive correlation between GABA and C-S synapse number, we did not observe a reduction in GABA levels in the two HD models relative to WT (**Table 1**). This is consistent with previous reports that have used ^1^H-MRS in zQ175 and R6/2 mouse models (Tkác et al., 2007; Heikkinen et al., 2012). A potential explanation is that ANOVAs of the neurochemical concentrations are based on the average concentration across multiple mice within the same genotype and the variability within groups overwhelmed the group differences across genotypes. However, the correlation analyses are conducted by plotting individual concentrations for each analyzed animal and their corresponding C-S synapse number. This suggests that individual levels of GABA are inherently associated with the C-S synapse density regardless of the genotype.

**Figure 4.**
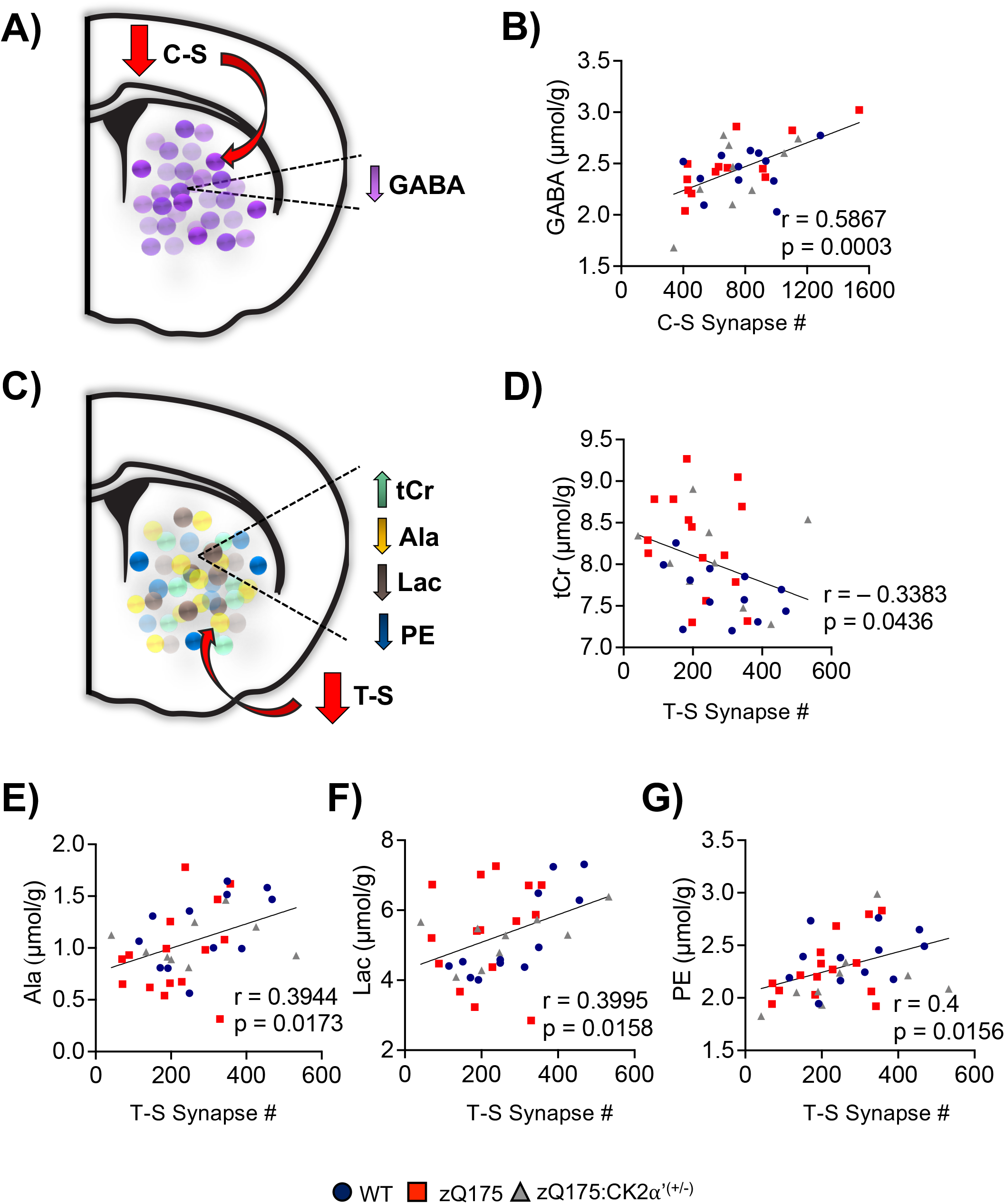
Selective neurochemical alterations differentially correlate with striatal synaptic circuits. **A**) Illustration of C-S correlation with metabolites (GABA). **B**) Correlation analysis of C-S synapse density vs. GABA where raw synapse numbers are plotted against corresponding neurochemical levels. **C**) Illustration T-S correlation with metabolites. **D-G**) Correlation analysis of neurochemicals vs T-S synapse density. r value represents Pearson’s correlation. Data from WT n=12, zQ175 n=15, zQ175:CK2α’^(+/−)^ n=9 corresponding to all three time points (3, 6 and 12 months) is shown.

In T-S circuitry, we found Ala, Lac, PE had positive correlations of a meaningful magnitude (**Table 3, Fig 4C-G**). tCr and tNAA/tCr had slightly weaker correlation values. Conservative multiple testing adjustment of p-values across all the metabolites did not maintain their statistical significance despite the global effect being significant. This could be due to the conservativism of the multiple testing adjustment, the high number of MRS measurements being tested (n=20), and the relatively small sample size which together may result in rejecting significant associations that in reality are meaningful. Among the neurochemicals that showed significant correlations with synapse density based on unadjusted p-values, Lac concentration did not show a significant alteration across the different genotypes and Ala was significantly altered only at 6 months between WT and zQ175 (**Table 1**). However, both Lac and Ala presented a trend towards lower levels in zQ175 compared with WT mice and were partially rescued in zQ175:CK2α’^(+/−)^ (**Table 1**). Similar trends for Ala and Lac were shown in symptomatic R6/2 mice (Tkác et al., 2007). PE was significantly lower in both zQ175 and zQ175:CK2α’^(+/−)^ than WT at 6 months (**Fig. 2C, Table 1**). tCr showed an increasing trend over time in both zQ175 and zQ175:CK2α’^(+/−)^ and was higher than WT at 6 and 12 months (**Fig. 2F**). These data indicate that by combining the levels of GABA, Ala, Lac, PE, and tCr, it could be possible to evaluate circuit-dependent synapse content during HD.

## Discussion

In this first direct assessment of the longitudinal association between neurochemical abnormalities and excitatory synapse density in the mouse brain we show that distinct neurochemical levels significantly correlate with different striatal glutamatergic synaptic input pathways in the zQ175 model, suggesting that ^1^H-MRS may distinguish circuit-dependent synapse changes in HD.

The current immunofluorescence data confirmed and expanded previous observations regarding changes in T-S and C-S synapse density in zQ175 and other mouse models (Raymond et al., 2011; Deng et al., 2014; Mckinstry et al., 2014; Gomez-Pastor et al., 2017) and showed that C-S synapse depletion occurs at advanced disease stages in fully symptomatic animals, while T-S synapse depletion occurs earlier. We also showed that reducing levels of CK2α’, a kinase previously associated with the dysregulation of synaptic activity in HD (Gomez-Pastor et al., 2017; Yu et al., 2020), prevented loss of T-S synapses without altering C-S synapse density, supporting previous data in younger mice (Gomez-Pastor et al., 2017). In older mice (12 months), reduction of CK2α’ did not completely suppress T-S synapse loss but rather delayed its onset. It is unknown how CK2α’ haploinsufficiency selectively improved T-S synapse loss during early stages of disease, although this could be related to a different regulatory role of CK2α’ between D1 and D2-MSNs, the latter being preferentially altered in HD (Reiner et al., 1988; Rebholz et al., 2013). Further studies are warranted to uncover the specific role of CK2α’ in the differential regulation of T-S and C-S synapses.

^1^H-MRS in patients with SCAs with different degrees of synapse loss showed a similar ranking in the severity of neurochemical alterations suggesting that neurochemical abnormalities across different SCAs, and perhaps other neurodegenerative diseases, may reflect abnormalities in synaptic function or density (Öz et al., 2011b; Joers et al., 2018). Immunohistochemical analyses for synaptic vesicles (SV2A) combined with antemortem ^1^H-MRS in brains from patients with Alzheimer’s disease (AD) showed decreased tNAA/tCr was associated with loss of synapses and early Tau pathology while increased Ins/tCr was associated with the occurrence of amyloid plaques in AD (Murray et al., 2014). Furthermore, a recent study in an AD mouse model with engrafted WT neural stem cells in the hippocampus showed changes in NAA and Glu that paralleled altered expression of synaptic proteins like PSD-95 and synaptophysin, and number of synapses (measured by electron microscopy) (Zhang et al., 2017). In a different study, ^1^H-MRS and synaptic protein analyses by immunoblotting in mice fed with a high fat diet showed increases in tCr and Gln that paralleled decreased synaptic proteins PSD-95 and VGlut1 (Lizarbe et al., 2018). Although correlations between neurochemicals and synapse numbers were not investigated with data collected in the same brains at the same time point/age in these prior studies, these reports established a plausible connection between key neurochemical alterations and synapse density.

In HD, ^1^H-MRS analyses have also highlighted tNAA, Ins and tCr as biomarkers of neurodegeneration (Tkác et al., 2007; Sturrock et al., 2010, 2015; Heikkinen et al., 2012; Peng et al., 2016). However, the differences between HD mouse models and patients regarding alterations of these specific neurochemicals, necessitate a careful analysis and validation of their potential relevance in defining synapse loss. In pre-symptomatic patients with HD, tNAA is lower than unaffected individuals and decreases further during disease progression, correlating with impaired motor and cognitive function (Leavitt et al., 2011). However, changes in tNAA are mostly observed only in symptomatic HD mice. Similarly, increased Ins was only observed in symptomatic HD mice (Tkác et al., 2007; Heikkinen et al., 2012; Yu et al., 2020). Therefore, tNAA and Ins seem to represent biomarkers for advanced disease in HD mouse models. This is supported by longitudinal MRS performed at 7T in zQ175, showing no significant alterations in tNAA or Ins up to 12 months (Heikkinen et al., 2012). Our current data at 9.4T validated these results for tNAA up to 12 months, but we previously reported changes in older (22 months) zQ175 mice (Yu et al., 2020). In addition, we found higher Ins in zQ175 vs. WT mice, which was significant at 6 months. Discrepancies between HD mouse models and patients were also reported for tCr (Sánchez-Pernaute et al., 1999; Sturrock et al., 2015; Adanyeguh et al., 2018). We confirmed an age-dependent increase in tCr in zQ175 and reported additional alterations in Gln, tCho and tNAA/tCr between WT and zQ175. These changes were not attributed to ventricular enlargement or large brain volume alterations between zQ175 and WT mice since no differences in these parameters were observed even in older mice (Yu et al., 2020). Also, if CSF contribution to the MRS voxel was significantly higher in HD vs. WT mice, we would expect a lowering of all metabolite levels in HD vs. WT, however some metabolites are higher (e.g. tCr, Gln, Ins) while others (tCho, PE, Ala) are lower in HD vs. WT. We conclude that Gln, tCr, tCho and tNAA/tCr represent biomarkers that monitor disease progression in zQ175 mice better than tNAA or Ins.

Among all tested neurochemicals, we only detected a significant alteration in Gln when comparing zQ175 and zQ175:CK2α’^(+/−)^. This was unexpected due to this latest model showing improvements in several HD-like phenotypes including decreased HTT aggregation and astrogliosis, increased synaptic density and neuronal excitability, and improved motor coordination (Gomez-Pastor et al., 2017; Yu et al., 2020). Gln levels increased over time in all mice but were significantly higher in zQ175 compared to WT starting at 3 months, consistent with a previously reported excitotoxic state (Fan and Raymond, 2007; Hassel et al., 2008). Analyses in R6/2 mice showed early increases in Gln levels (Tkác et al., 2007). Increased Gln has been associated with an imbalance in Glu–Gln cycling between neurons and astrocytes, reflecting compromised glutamatergic neurotransmission (Liévens et al., 2001; Behrens et al., 2002). Notably, no significant difference in Gln levels were found between WT and zQ175:CK2α’^(+/−)^. Therefore, it is reasonable to hypothesize that early decreased Gln in zQ175:CK2α’^(+/−)^ vs. zQ175 could be associated with improved neuronal excitability. In support of this hypothesis, zQ175:CK2α’(+/−) previously showed improved AMPA-mediated excitatory transmission in the dorsolateral striatum when compared to zQ175 (Yu et al., 2020).

Correlation analyses revealed a direct association between changes in specific neurochemicals and circuit-dependent synapse changes. We found a significant positive correlation between levels of GABA and C-S synapse density. Deficiency in GABA signaling was associated with HD and other movement disorders (Hsu et al., 2018). Lower GABA content in the dorsal striatum and cortex was reported in postmortem brain from patients with HD (Spokes et al., 1980). Metabolic profiling using ^13^C labeling and mass spectrometry in symptomatic R6/2 mice also showed decreased GABA synthesis (Skotte et al., 2018). However, ^1^H-MRS did not reveal changes in GABA levels in R6/2 compared with WT (Tkác et al., 2007). We also did not detect significant differences in GABA between WT and zQ175. The discrepancy between the lack of overall alterations in GABA and its correlation with C-S synapse density could be explained by individual levels of GABA being inherently associated with C-S synapse density regardless of genotype. On the other hand, we observed significant correlations between T-S synapse density and Ala, Lac, PE, and tCr. Ala and other neutral amino acids are low in plasma from patients with HD, which correlates with symptom status (Reilmann et al., 1995; Underwood et al., 2006). Lac is an end-product of glycolysis associated with glutamatergic synaptic activity (Solís-Maldonado et al., 2018). ^1^H-MRS Lac changes in patients with HD is controversial (Adanyeguh et al., 2018) but studies using ^13^C-Lac showed decreased Lac uptake in HD cells (Solís-Maldonado et al., 2018). Although not statistically significant, we observed an overall decrease of Ala and Lac in zQ175 relative to WT that was partially rescued in zQ175:CK2α’^(+/−)^. Alterations in PE and tCr, which are connected with neuronal integrity and function, have been consistently reported in mouse models and patients with HD and are associated with disease burden (Tkác et al., 2007; Sturrock et al., 2010, 2015; Adanyeguh et al., 2018). tCr increased over time in zQ175 and zQ175:CK2α’^(+/−)^ compared with WT and significantly correlated with T-S synapse density. Our data suggest that a combination of the levels of GABA, Ala, Lac, PE, and tCr could be used to assess levels of striatal glutamatergic synapses in HD.

Overall, this study demonstrates the feasibility of using ^1^H-MRS to potentially monitor synaptic changes *in vivo* during HD progression. A limitation of our study is the difference between the MRS volume, and the area covered in synapse density analyses. The correlation analyses could be improved in future studies by reducing the MRS VOI size and distributing the tissue sampling areas for immunohistochemistry further within the MRS VOI. In addition, while significant correlations between neurochemicals and synapse loss were found, we cannot discard these associations as being influenced by other neuropathological features. Therefore, further studies are needed to corroborate these findings in alternative models of HD and other neurodegenerative diseases to evaluate specific neurochemical biomarkers as a tool to monitor synaptic changes in future pre-clinical trials with HD models.

## Acknowledgements

We thank Dr. Dinesh Deelchand for guidance in data analysis and assistance in LCModel basis set generation.

## Authors’ Roles

R.G.P. and G. O. obtained funding for this study and designed the experiments. N.Z., K.G., J.C., and D.Y. performed the experiments. N.Z., K.G., J.C., and D.Y. prepared and analyzed the data. G.O. supervised the MRS data acquisition and analysis. R.G.P. supervised the *ex vivo* tissue collection and synapse analyses. L.E. supervised the statistical analyses. N.Z. and R.G.P. wrote the first draft of the manuscript and all authors edited subsequent versions and approved the final version of the manuscript.

## Financial Disclosure

Nothing to report.

## Notes

**Relevant conflicts of interest/Financial Disclosure:** Nothing to report

**Funding Sources:** This work was supported by the University of Minnesota Biomedical Research Awards for Interdisciplinary New Science (to R.G.P and G.O) and the National Institute of Health NINDS (R01 NS110694) (to R.G.P). The Center for Magnetic Resonance Research is supported by the National Institute of Biomedical Imaging and Bioengineering (NIBIB) grant P41 EB027061, the Institutional Center Cores for Advanced Neuroimaging award P30 NS076408 and the W.M. Keck Foundation.

### Competing Interest Statement

The authors have declared no competing interest.

### Summary of Updates

This version of the manuscript contains a new Figure and a revised results section

